# Methyl-TWAS: A powerful method for *in silico* transcriptome-wide association studies (TWAS) using long-range DNA methylation

**DOI:** 10.1101/2023.11.10.566586

**Authors:** Soyeon Kim, Yidi Qin, Hyun Jung Park, Molin Yue, Zhongli Xu, Erick Forno, Wei Chen, Juan C. Celedón

**Author notes:** Shared first authors (Soyeon Kim, Yidi Qin). Shared senior authors (Soyeon Kim, Juan C. Celedón). **Correspondence:** Juan C. Celedón, MD, DrPH Division of Pulmonary Medicine UPMC Children’s Hospital of Pittsburgh 4401 Penn Avenue, Pittsburgh, PA 15224.

## Abstract

*In silico* transcriptome-wide association studies (TWAS) are commonly used to test whether expression of specific genes is linked to a complex trait. However, genotype-based *in silico* TWAS such as PrediXcan, exhibit low prediction accuracy for a majority of genes because genotypic data lack tissue- and disease-specificity and are not affected by the environment. Because methylation is tissue-specific and, like gene expression, can be modified by environment or disease status, methylation should predict gene expression with more accuracy than SNPs. Therefore, we propose Methyl-TWAS, the first approach that utilizes long-range methylation markers to impute gene expression for *in silico* TWAS through penalized regression. Methyl-TWAS 1) predicts epigenetically regulated/associated expression (eGReX), which incorporates tissue-specific expression and both genetically- (GReX) and environmentally-regulated expression to identify differentially expressed genes (DEGs) that could not be identified by genotype-based methods; and 2) incorporates both *cis-* and *trans-* CpGs, including various regulatory regions to identify DEGs that would be missed using *cis*-methylation only. Methyl-TWAS outperforms PrediXcan and two other methods in imputing gene expression in the nasal epithelium, particularly for immunity-related genes and DEGs in atopic asthma. Methyl-TWAS identified 3,681 (85.2%) of the 4,316 DEGs identified in a previous TWAS of atopic asthma using measured expression, while PrediXcan could not identify any gene. Methyl-TWAS also outperforms PrediXcan for expression imputation as well as *in silico* TWAS in white blood cells. Methyl-TWAS is a valuable tool for *in silico* TWAS, leveraging a growing body of publicly available genome-wide DNA methylation data for a variety of human tissues.

## INTRODUCTION

Transcriptome-wide association studies (TWAS) are commonly used to test whether expression of specific genes is linked to a complex trait. When gene expression is not directly measured but genome-wide genotypic data *<u>or</u>* summary statistics from genome-wide association studies (GWAS) or an expression quantitative trait locus (eQTL) study are available, gene expression can be imputed to conduct *in silico* TWAS.

Methods used for imputation of gene expression to conduct *in silico* TWAS include PrediXcan, which employs SNPs within 1 Mb from gene regions^1^; S-PredXcan^2^, which incorporates summary statistics from GWAS; and BGW-TWAS^3^ and OTTERS^4^, which employ summary statistics from eQTL studies. While commonly used for *in silico* TWAS, these genotype-based approaches have low average prediction accuracy for gene expression (e.g., Test R^2^=0.02-0.05 for PrediXcan ^1,4^; 0.04 for FUSION; 0.05 for PRS-CS^4^), likely because SNPs do not account for the tissue specificity of gene expression (Figure 1A) *<u>or</u>* modification of gene expression by epigenetic mechanisms^5^, environmental factors^6^ or disease status. When prediction accuracy for gene expression is low, *in silico* TWAS yield erroneous results^7^.

**Figure 1.**
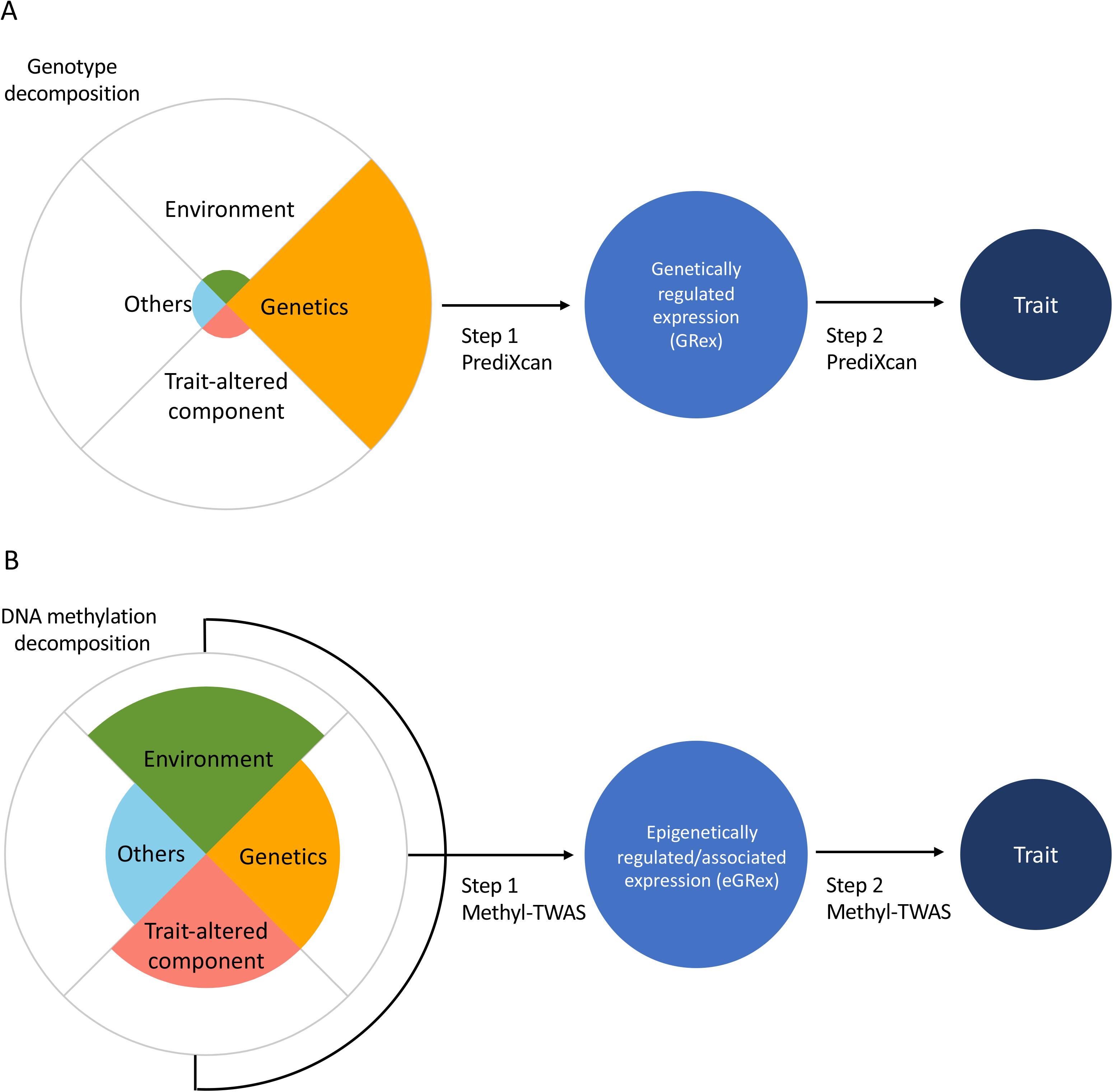
Comparison of conceptual components of genotypes vs. DNA methylation. A. PrediXcan uses SNPs to impute gene expression, thus not accounting for tissue specificity or non-genetic regulation of gene expression (i.e., by environment or disease status). B. Methyl-TWAS incorporates genetic regulation, tissue specificity, and modification of DNA methylation and gene expression by environment or disease status when imputing gene expression levels, which refers to epigenetically regulated or associated expression (eGRex).

Because DNA methylation is tissue-specific^8^ and, like gene expression, can be modified by environment or disease status (Figure 1B), methylation markers such as CpGs can predict gene expression with more accuracy than SNPs. Indeed, expression quantitative trait methylation (eQTM) studies have identified numerous associations between CpGs and gene expression in skeletal muscle^9^, blood^10^, and nine separate tissues in GTex data^11^.

We recently identified 2.1 million eQTM loci in nasal epithelium of youth with and without atopic asthma (both in *cis-* and *trans-^12^*) and found that several gene regulatory elements are enriched in eQTM CpGs. Further, we showed that *cis*-eQTM CpGs are enriched in the enhancer region of their associated genes^13^, that *trans*-eQTM CpGs are enriched near transcription factors and miRNAs, and that these CpGs are significantly associated with their target genes*^12^*. This suggests that DNA methylation may regulate gene expression by affecting gene regulatory elements.

In this work, we propose the first methylation-based *in silico* TWAS method, referred to as Methyl-TWAS. Methyl-TWAS: 1) predicts epigenetically regulated/associated expression (eGReX), which incorporates genetically- (GReX), and environmentally-regulated expression, trait-altered expression, and tissue-specific expression to identify differentially expressed genes (DEGs) that could not be identified by genotype-based methods (**Figure 1B**); and 2) incorporates both *cis-* and *trans-* CpGs, including enhancers, promoters, transcription factors, and miRNA regions to identify DEGs that would be missed using *cis*-DNA methylation-based methods. To achieve this, we extend our previous method, geneEXPLORE^14^, which incorporates long-range DNA methylation (10 Mb away) from promoter regions for gene expression imputation, to identify DEGs using *in silico* TWAS. Further, we compare a DNA methylation-based method (Methyl-TWAS) with a genotype-based method (PrediXcan) in predicting gene expression levels and yielding accurate results from *in silico* TWAS using imputed expression levels. For this, we use our in-house data consisting of genome-wide genotypes and RNA-seq data from nasal epithelial samples (n=469), and DNA methylation and RNA-seq data from nasal epithelial samples (n=455) collected in individuals with and without asthma aged 9 to 20 years who participated in the Epigenetic Variation and Childhood Asthma in Puerto Ricans study (EVA-PR).

We compare the gene expression prediction performance of Methyl-TWAS with that of three existing methods: PrediXcan, Hong and Rhee, and DNA combi. For gene expression imputation, PrediXcan utilizes *cis*-SNPs (within 1 Mb from promoter regions); the Hong and Rhee approach^15^ utilizes *cis*-CpGs; and DNA combi incorporates *cis*-genotypes and *cis*-CpGs. Because only PrediXcan has been used for *in silico* TWAS, we compare Methyl-TWAS with PrediXcan for accuracy of the results of an *in silico* TWAS versus those of a TWAS using measured gene expression in nasal epithelium. Further, we demonstrate the use of Methyl-TWAS in another tissue, white blood cells, in EVA-PR.

## SUBJECTS AND METHODS

### Study population and procedures

Subject recruitment and the study protocol for the Epigenetic Variation and Childhood Asthma in Puerto Ricans study (EVA-PR) have been previously reported^48^. In brief, EVA-PR is a case-control study of asthma in individuals aged 9 to 20 years from Puerto Rico. Participants with and without asthma were selected from households in San Juan (PR) between February 2014, and May 2017 using a multistage probability sampling approach. Of 638 households with eligible subjects, 543 (85.1%) youth (one per household) agreed to participate. There were no significant differences in age or sex between eligible subjects who did and did not agree to participate. The study protocol included questionnaires on respiratory health, collection of blood samples, and collection of nasal samples for DNA and RNA extraction. Cases had asthma, defined as physician-diagnosed asthma and wheeze in the prior year. Control subjects had neither physician-diagnosed asthma nor wheeze in the prior year. Serum IgE levels to each of five common allergens (house dust mite, German cockroach, cat dander, dog dander, and mouse urinary protein) were measured using the UniCAP 100 system (Pharmacia & Upjohn, Kalamazoo, Mich), and atopy was defined as at least one positive (≥0.35 IU/mL) IgE to at least one of the allergens tested. EVA-PR was approved by the institutional review boards of the University of Puerto Rico (San Juan, PR) and the University of Pittsburgh (Pittsburgh, PA). Written parental consent and participant assent were obtained for individuals younger than 18, while consent was obtained from participants 18 years and older.

### Genome-wide genotyping, RNA sequencing, and genome-wide study of DNA methylation of EVA-PR

Genome-wide genotyping was conducted using the HumanOmni2.5 BeadChip platform (Illumina Inc., San Diego, CA, USA), as previously reported^49^. Genotype imputation was performed with the Michigan Imputation Server ^17^, using the Haplotype Reference Consortium (HRC) r1.1 2016 ^18^ as the reference panel.

The HumanMethylation450 BeadChip (Illumina, Inc.) was used to measure genome-wide DNA methylation. Beta-values, which range from 0 to 1, were utilized to determine the percentage of methylation. To ensure compatibility with linear regression models that assume a normal distribution, we transformed the beta values into M-values using the formula M = log2(beta/(1-beta)). M-values provide a closer approximation to a normal distribution.

RNA-seq was conducted using the Illumina NextSeq 500 platform (Illumina, Inc.), with the resulting data expressed as transcripts per kilobase million (TPM) values. Genes with low expression levels (mean TPM < 1) were excluded from the analysis, as well as those lacking information on the transcription start sites in the UCSC genome browser. To facilitate data analysis, the TPM values were transformed to log2(TPM+1).

To account for potential variations caused by different cell types, a specific protocol was implemented in a subset of nasal samples (n=31). This involved selecting CD326-positive (cell-sorted) nasal epithelial cells before conducting DNA and RNA extraction. This procedure, along with other quality control measures, was previously described^48^.

After quality control, we had 469 subjects (228 cases and 241 controls) in whom both genotypes and RNA-seq data were available for this analysis, and 455 subjects (219 cases and 236 controls) in whom both DNA methylation and RNA-seq data were available for this analysis.

### Estimating epigenetically regulated/associated expression (eGReX)

In Stage I of Methyl-TWAS, we incorporated elastic-net^50^, a penalized linear regression method, to estimate the epigenetically regulated portion of gene expression (**Figure 1B**). The elastic-net method is widely used for genomic data analysis because it can handle correlated features in omics data. For each gene, we built a linear regression model to predict expression level using long-range CpGs.

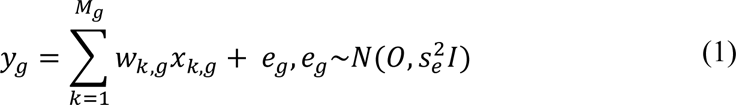

where *y*_*g*_ is the vector of expression level of gene *g*, *x*_*k*,*g*_ is the vector of DNA methylation M value at *k*-th methylation CpG for gene *g, M*_*g*_ is the number of methylation CpGs for the gene, *w*_*k*,*g*_ is the regression coefficient of the methylation CpGs. Here, we consider CpGs within 10 Mb from the promoter region. We chose ±10 Mb based on our previous research in breast cancer: we investigated all ranges from 1 Mb to the entire chromosome where the gene is located and found that after 10Mb, prediction accuracy is not much improved but computational time is dramatically increased^14^. Methyl-TWAS estimates the effect size through the elastic-net method where 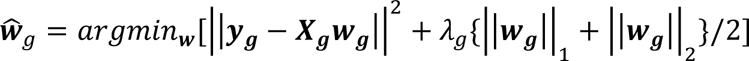 is the vector of 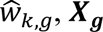 is the matrix of *x*_*k*,*g*_, and *λ*_*g*_ is the regularization parameter for gene *g*. ||***w***_***g***_|| is L1 penalty (the sum of absolute numbers) and ||***w***_***g***_|| is L2 penalty (the sum of squared numbers). The penalty, *λ*_*g*_, was selected through cross-validation using the R package glmnet.

We chose the elastic net method because, without any statistical tests, elastic-net automatically selects important variables that are associated with a response by making the multiple of the coefficients exactly zero due to the L1 penalty. Through this characteristic of inelastic-net, variables with zero coefficients are not selected in the model, and the variables with non-zero coefficients are selected in the model. By utilizing elastic-net, the model automatically selects CpGs associated with gene expression from tens of thousands of CpGs. Moreover, elastic-net works well in highly correlated datasets. Due to the L2 penalty in the model, elastic-net can select highly correlated variables together^50^. Since multiple methylation values are highly correlated due to biological interactions (e.g. SNPs in high linkage disequilibrium and CpGs in CpG islands) ^50-52^, the method has been used to analyze several types of omics data^50-52^.

In stage II, we use the estimated effect sizes 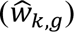 from Stage I to impute gene expression in a test data set (eGReX), 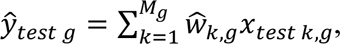 and then test for association between eGReX and phenotype, which is called TWAS using an empirical Bayesian linear regression framework in R package limma^53^.

### Method performance comparison

We compared the prediction performance of four methods: Methyl-TWAS, PrediXcan^1^, Hong and Rhee^15^, and DNA combi. We used tenfold cross-validation (CV) to measure CV-Pearson’s correlation (CV-r). Nine folds of data consisting of CpGs and/or SNPs, as well as gene expression (in terms of the number of samples) were used to build a model as a training data set. The model used methylation values and/or SNPs in the remaining fold (a validation data set) to predict gene expression. We repeated the procedure ten times until gene expression levels were imputed for all samples. We selected the model (or the penalty) that maximizes prediction accuracy using CpGs within 10 Mb from promoter regions. Prediction accuracy was measured by Pearson’s correlation coefficient between predicted and observed gene expression.

For better comparison across methods, we defined three sets of genes and investigated the prediction accuracy of gene expressions within each set: 1) Valid genes: a set of genes whose Pearson’s correlation is higher than 0.1 using at least one of four methods; 2) Immune genes: a gene list defined by taking genes within 17 categories from IMMPORT (Bioinformatics For the Future of Immunology) database ^22^; 3) DEGs: differentially expressed genes at FDR-P < 0.01 from TWAS analysis using observed gene expression from EVA-PR data (previously reported ^20^). Within each gene set, we summarized and compared the prediction accuracy of each method by calculating the mean and median values of Pearson’s correlation derived from cross-validation (CV-r).

To identify which genes can be better predicted by Methyl-TWAS or PrediXcan, we conducted two-sided z tests for two correlations to assess whether the difference between CV-r using Methyl-TWAS and CV-r using PrediXcan is significant for each gene. For a gene with correlation (r) from method 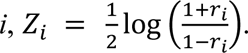 The standard deviation of the difference is 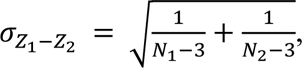 and the test statistic is given by 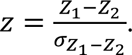 After the test, we conducted gene ontology (GO) enrichment analysis using the genes better predicted by each method (CV-r > 0.5, Bonferroni adjusted P < 0.05) ^54,55,56^.

For immunity-related gene sets, we compared the prediction accuracy of the four methods against such accuracy for the same number of randomly selected genes from non-immune gene sets. Moreover, for methyl-TWAS only, we compared CV-Pearson’s correlations of eight immune gene lists (pathways), which have at least 30 genes. The immune gene lists were gotten from IMMPORT.

For DEG sets, we used the same approach to compare the prediction accuracy of the four methods for DEGs against such accuracy for the same number of randomly selected non-DEGs. One-sided t-tests were used to test if the mean of CV-r of immune genes or that of DEGs is higher than the randomly selected genes.

### *In silico* TWAS of atopic asthma in nasal epithelium

For the *in-silico* TWAS of atopic asthma in EVA-PR, we conducted nested cross-validation to minimize overfitting. Data were divided into a training set (4/5 of the samples) and a test set (1/5 of the samples). Using the training dataset, 10 fold CV were conducted (therefore, 4/50 of samples are in each fold) to select a model that minimizes mean squared error. We predicted gene expression of the test dataset using the estimated coefficients from the trained model, and DNA methylation or SNPs in the test dataset as the input using Methyl-TWAS or PrediXcan respectively. We repeated the procedure five times until we predicted gene expression levels for all samples.

Using predicted gene expression of EVA-PR through nested CV, we conducted differential gene expression analysis. DEGs were identified through an empirical Bayesian linear regression framework in R package limma ^53^. The multiple regression model was adjusted for age, sex, RNA-seq plate, sample sorting protocol (whether CD326-positive or not), and the top five principal components (PCs) derived from genotypic data. We identified DEGs at a false discovery rate adjusted P value (FDR-P) <0.01.

### Testing in an independent cohort (GSE65205)

We used the public dataset provided by Yang et al.^33^ (GSE65205) as test data to show the prediction accuracy of the models trained using EVA-PR data. GSE65205 consists of methylation 450K array data and gene expression array data (G3 SurePrint 8x60k, Agilent Technologies) in nasal epithelium from 69 predominantly African American children (36 with atopic asthma and 33 controls). Methyl-TWAS models were built using EVA-PR for each gene, and models with a minimum 10-fold CV error were selected. Using the CpGs from the test dataset (GSE65205) as input for the models, we predicted expression levels for 10,477 genes. These genes were among 12,745 valid genes and also available in the microarray data set. Test-Pearson’s correlation coefficient between the predicted and observed gene expression (microarray log2 normalized value) of the test dataset was then calculated. Differential gene expression analysis was conducted using predicted gene expression of GSE65205 data as well. Due to limited phenotypic information, we only adjusted the model for age and sex. Due to small sample size, we used FDR-P < 0.05 to identify DEGs. In the TWAS Manhattan plot, Bonferroni P <0.05 was used to label genes passing the threshold.

### Cross-validation in white blood cell (WBC) data from EVA-PR participants

Of the 543 EVA-PR participants, 464 had genome-wide methylation data for WBCs. To compare the prediction performances of Methyl-TWAS and PrediXcan using WBC data, we performed a 10-fold CV to measure CV-Pearson’s correlation in the same way as we did for nasal epithelium. To train the model, Methyl-TWAS used CpGs as predictors and gene expression as the outcome using WBC data, while PrediXcan used non-tissue-specific SNPs as predictors and gene expression in WBCs as the outcome. We further performed TWASs using gene expression predicted by Methyl-TWAS and PrediXcan. In addition to atopic asthma status, the TWAS model adjusted for age, sex, and the top five PCs derived from genotypic data. We identified DEGs at FDR-P <0.05.

## RESULTS

### Methods overview

The aim of Methyl-TWAS is to identify DEGs that affect a trait of interest through *cis- and trans-* DNA methylation. In step 1, Methyl-TWAS first estimates weights for long-range (*cis-* and *trans-* <10 Mb) CpGs per gene (*g)* using the elastic-net penalized regression models on a training dataset (**Figure 2**). In this step, Methyl-TWAS drops many CpGs not associated with the gene out of the model because of L1 penalty in elastic-net regression and estimates higher weights for more impactful CpGs that regulate or are associated with expression of each gene. In step 2, Methyl-TWAS predicts eGReX using the coefficients (weights) estimated in the first step and the selected CpGs in a test DNA methylation dataset. In step 3, using the imputed gene expression from step 2, Methyl-TWAS conducts *in silico* TWAS to identify DEGs in a trait or condition.

**Figure 2.**
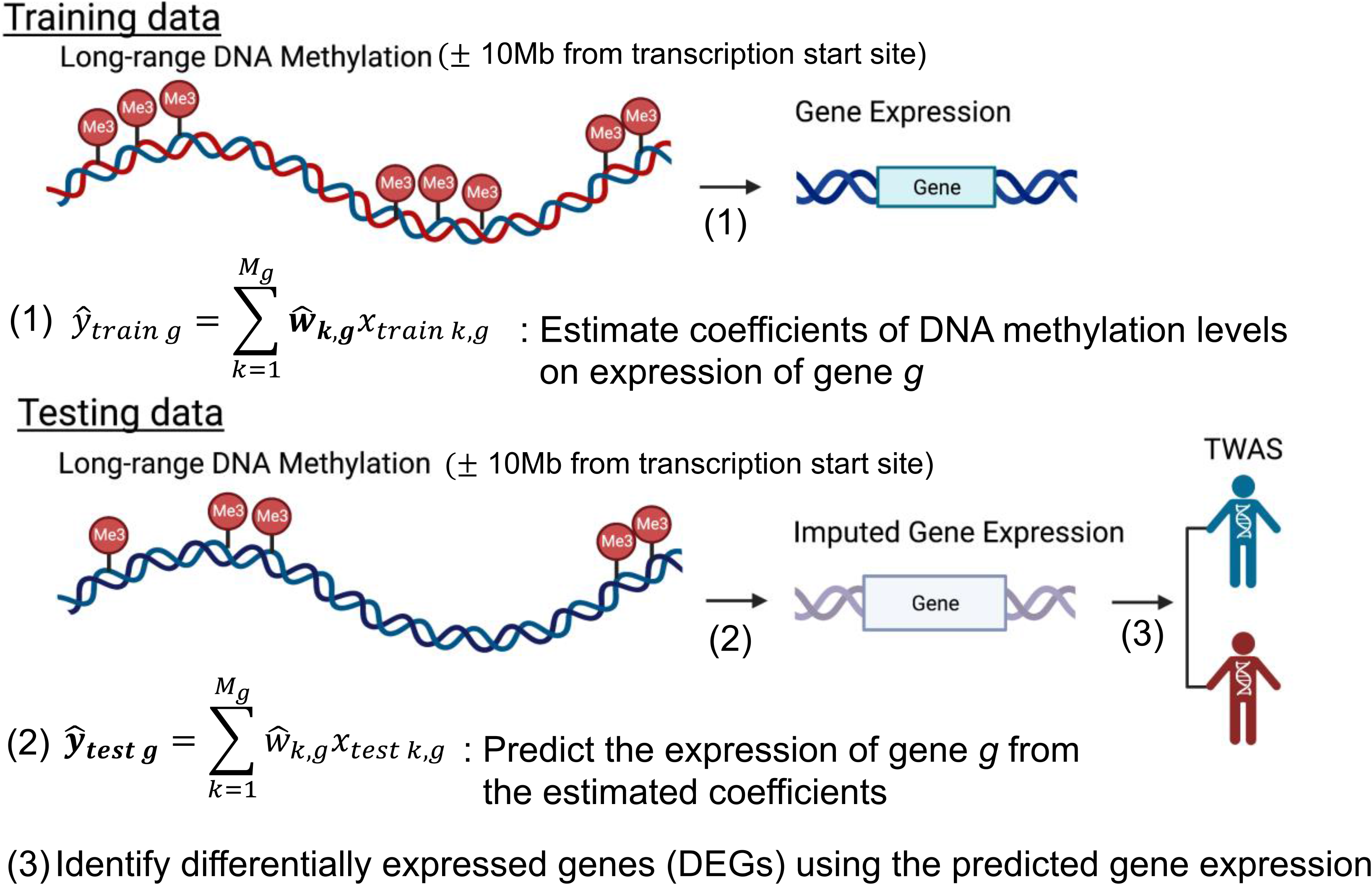
Methyl-TWAS framework. Methyl-TWAS (1) estimates weights for long-range DNA methylation markers from the elastic-net penalized regression models using the training data set, (2) predicts gene expression using a test data set, and (3) identifies differentially expressed genes or DEGs using the imputed gene expression levels for *in silico* TWAS

We have made the R-package, Methyl-TWAS, publicly available at https://github.com/YidiQin/MethylTWAS. The package offers two modes. In the first mode, users can train the model and conduct TWAS using any tissue. Users input the training (reference) data consisting of DNA methylation and gene expression data for the same subjects to estimate tissue-specific effects of DNA methylation on expression for each gene and then input testing data of DNA methylation and a phenotype to impute gene expression and then conduct *in silico* TWAS. In the second mode, users can skip step 1 and just conduct TWAS using a test data set (DNA methylation) of nasal (airway) epithelium. Users input DNA methylation and a phenotype and then, Methyl-TWAS uses the estimated coefficients (calculated using our reference data from EVA-PR and uploaded to GitHub) to impute gene expression and conduct *in silico* TWAS without requiring further reference data.

### Methyl-TWAS outperforms other methods, especially PrediXcan, in predicting gene expression in nasal epithelium

Using genome-wide SNPs and CpGs and RNA-seq data from nasal epithelial samples of EVA-PR participants, we compared the prediction accuracy for gene expression of Methyl-TWAS, PrediXcan^1^, the Hong and Rhee approach^15^, and DNA combi. We measured CV-r between predicted and observed gene expression for 14,260 genes. Of these, 12,745 genes were “valid” genes with r >0.1 (corresponding to R^2^ >0.01) using at least one method, as suggested by various *in silico* TWAS methods^4,16,17^.

On average, Methyl-TWAS outperforms the other methods (**Figure 3, panel A**) (mean r =0.252, 0.034, 0.204, and 0.211 for Methyl-TWAS, prediXcan, Hong and Rhee, and DNA combi respectively). The prediction accuracy of prediXcan is the worst (**Figure 3, panels A, B,** an**d D**), with Methyl-TWAS predicting about five times more expression levels better than PrediXcan (10,578 vs. 2,167). Adding SNPs to *cis-* methylation data (DNA combi) does not improve prediction compared with using methylation data only in the same range (1M) (average r=0.21 vs 0.20 on average, **Figure 3A**). In fact, *cis-*DNA methylation (Hong and Rhee) predicts the expression of more genes than DNA combi (**Figure 3C**) (7,938 vs. 6,333). On the other hand, adding CpGs to SNPs (DNA combi) does significantly improve prediction accuracy compared with using SNPs only (PrediXcan) (**Figure 3, panels A** and **D**).

**Figure 3.**
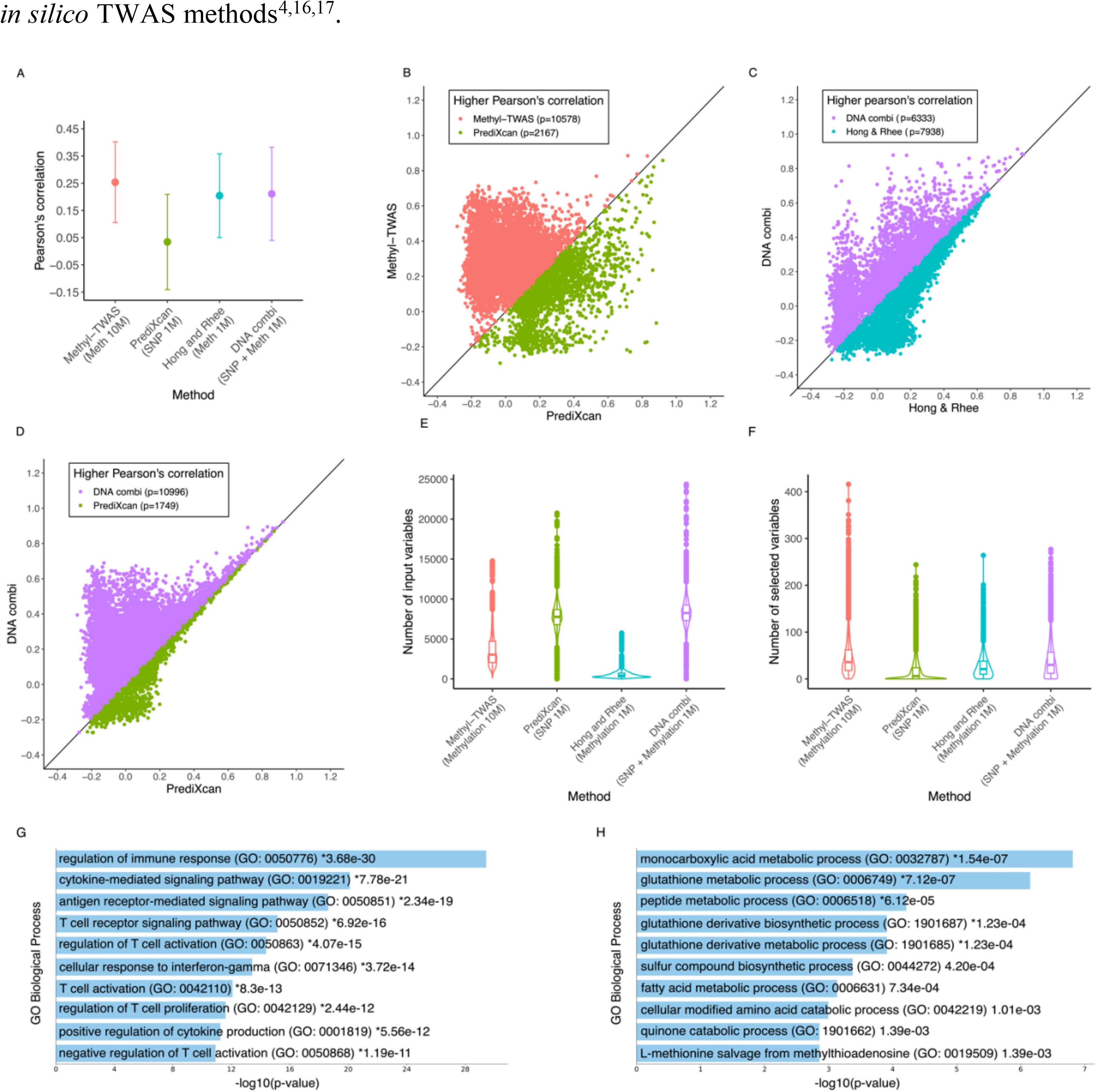
Comparison of prediction accuracy using Methyl-TWAS (CpGs ±10Mb from the promoter region), PrediXcan (SNP ±1 Mb from the promoter region), Hong and Rhee (CpGs ±1 Mb from the promoter region), and DNA combi (SNPs and CpGs ±1 Mb from the promoter region) in EVA-PR data. A. Pearson’s correlation (r) between observed and predicted gene expression using 10-fold cross-validation (CV) of EVA-PR data. We included 12,745 genes for which at least one method predicted gene expression with CV-r > 0.1. Mean and a standard deviation bar. B. Comparison of prediction accuracy between PrediXcan and Methyl-TWAS. C. Comparison of prediction accuracy between Hong and Rhee and DNA combi. D. Comparison of prediction accuracy between PrediXcan and DNA combi. E. Comparison of input variables (the number (#) of SNPs and/or # of DNA methylation CpGs) for the 4 methods F. Comparison of the number of selected variables by the method (# of SNPs and/or # of DNA methylation CpGs) for the 4 methods. G. GO Pathway analysis using 561 genes significantly better predicted by Methyl-TWAS (CV-r>0.5 & FDR < 0.05). H. GO Pathway analysis using 239 genes significantly better predicted by PrediXcan (CV-r>0.5 & FDR < 0.05).

When we compare methods that only use DNA methylation, Methyl-TWAS outperforms Hong and Rhee’s approach, which only use *cis*-CpGs (**Figure 3, panel A**), supporting larger collective effects of long-range CpGs on nasal epithelial gene expression than those of *cis*-CpGs.

We also checked the number of input SNPs or CpGs and associated variables selected by elastic-net in the final model. Even though Methyl-TWAS incorporates fewer than half the input variables required by PrediXcan on average (p=3,622 vs. p=7,721) (**Figure 3, panel E**), the number of selected variables using Methyl-TWAS is about three times higher (s=45.6), on average, than that using PrediXcan (s=16.4)(**Figure 3, panel F**). This implies that even though there are more SNPs than CpGs, fewer SNPs affect gene expression than CpGs do. While adding *ci*s-SNPs to *cis*-CpGs does not make a difference for prediction accuracy, it is nearly additive with regard to the average number of selected variables: using *cis*-DNA methylation (Hong and Rhee)=26.8, cis-SNPs (PrediXcan)=16.4, and using both (DNA combi)= 40.0. Thus, adding selected (associated) SNPs to CpGs has minimal additional effects on imputing gene expression.

For genes with expression predicted with high accuracy (r>0.5), we compared the prediction accuracy of PrediXcan with that of Methyl-TWAS to test whether there are statistically significant differences for each gene, finding that expression of 239 genes is significantly better predicted by PrediXcan (**Table S1**), while that of 561 genes is significantly better predicted by Methyl-TWAS (**Table S2**). Interestingly, while genes whose expression is better predicted by PrediXcan are enriched in metabolic, catabolic, and biosynthetic pathways, those whose expression is better predicted by Methyl-TWAS are enriched in immune pathways including cytokine, antigen receptor, T-cell, and interferon-gamma (**Figure 3, panels G-H**). The top 30 genes whose expression was significantly better predicted by Methyl-TWAS (mostly r >=0.6) were poorly predicted by PrediXcan, showing negative correlations (**Table 1**). The top 3 genes were *NTRK2, NLRC3*, and *CST1*, all relevant to asthma*. NTRK2 (TrkB) NTRK2* expression is increased in the airways of subjects with asthma and may play a role in airway hyper-responsiveness^18^, *NLRC3* influences T cell regulation^19^, and *CST1* (the most significant DEG for atopic asthma in EVA-PR^20^) promotes eosinophilic airway inflammation in asthma^21^.

**Table 1.** 30 most significant genes that are better predicted by Methyl-TWAS than PrediXcan. CV-R is Pearson’s correlation coefficient between predicted gene expression and observed gene expression using cross-validation (CV). The genes are sorted by the z statistics showing the difference. All adjusted p-values are less than 10^-300^.

| Gene ID | CV-R using Methyl-TWAS | CV-R using PrediXcan | Test statistics, z |
| --- | --- | --- | --- |
| *NTRK2* | 0.71 | -0.16 | 84.29 |
| *NLRC3* | 0.67 | -0.2 | 81.3 |
| *CST1* | 0.69 | -0.17 | 80.8 |
| *THEMIS* | 0.66 | -0.2 | 79.76 |
| *SLAMF6* | 0.65 | -0.22 | 79.68 |
| *CD247* | 0.66 | -0.19 | 79.33 |
| *CST2* | 0.68 | -0.16 | 78.9 |
| *CPA3* | 0.71 | -0.1 | 78.54 |
| *CXCR6* | 0.69 | -0.14 | 78.32 |
| *PYHIN1* | 0.66 | -0.18 | 77.76 |
| *CDH26* | 0.66 | -0.17 | 77.54 |
| *CD3G* | 0.65 | -0.2 | 77.48 |
| *CD244* | 0.66 | -0.17 | 76.88 |
| *KCNJ16* | 0.69 | -0.1 | 76.23 |
| *MS4A6A* | 0.63 | -0.22 | 76.19 |
| *PRF1* | 0.63 | -0.2 | 75.52 |
| *FCGR3A* | 0.65 | -0.17 | 75.49 |
| *CD48* | 0.65 | -0.17 | 75.47 |
| *ABI3* | 0.62 | -0.2 | 75.03 |
| *TPSAB1* | 0.67 | -0.13 | 74.93 |
| *FETUB* | 0.67 | -0.13 | 74.87 |
| *TYROBP* | 0.61 | -0.22 | 74.67 |
| *GIMAP6* | 0.63 | -0.18 | 74.62 |
| *FASLG* | 0.6 | -0.24 | 74.36 |
| *CD3D* | 0.64 | -0.17 | 74.23 |
| *CD3E* | 0.62 | -0.19 | 73.75 |
| *CANT1* | 0.6 | -0.23 | 73.53 |
| *SCGB3A1* | 0.62 | -0.19 | 73.43 |
| *NKG7* | 0.59 | -0.23 | 73.13 |
| *C1QA* | 0.61 | -0.21 | 73.11 |

### Methyl-TWAS better predicts expression of immunity-related genes and DEGs in atopic asthma

To identify which types of genes are better predicted by Methyl-TWAS, we investigated immunity-related genes, as they are relevant to asthma. First, we compared the predictive accuracy of Methyl-TWAS for expression of immunity-related genes in nasal epithelium with that of other methods. We retrieved 753 immunity-related genes from IMMPORT (Bioinformatics For the Future of Immunology)^22^. Methyl-TWAS predicts the expression of these genes significantly better than that of the same number of randomly selected genes (P < 2.2×10^-16^) (**Figure 4, panel A**). Hong and Rhee’s approach and DNA combi also predict expression of these immunity-related genes better than that of random genes (P < 2.2×10^-16^ and P=9.6 x10^-11^, respectively), but PrediXcan does not (P=0.99). Further, Methyl-TWAS outperforms the other three methods in predicting expression of these genes. In particular, while CV-r for immunity-related genes using Methyl-TWAS is 0.32, that for PrediXcan is 0.01 on average. This shows that the estimated collective effects of CpGs on expression of immunity-related genes are greater than those of SNPs.

**Figure 4.**
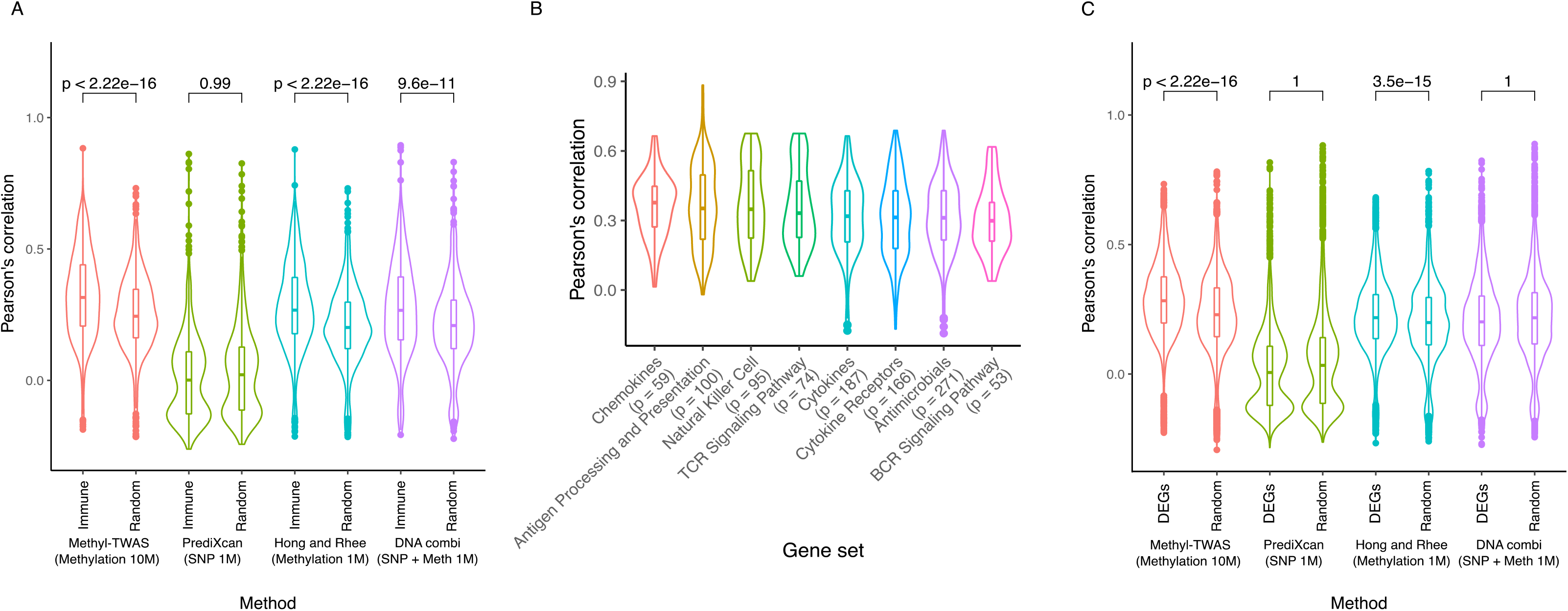
Prediction accuracy (CV-Pearson’s correlation, r) of expression levels for various categories of genes using four different methods. A. Accuracy of predicting expression of 753 immunity-related genes versus that of the same number of randomly selected genes using the four methods. The list of immunity-related genes was obtained from the IMMPORT database ^22^. B. Accuracy of predicting expression of immunity-related genes in 8 pathways using Methyl-TWAS. The pathways/gene lists were obtained from the IMMPORT database ^22^. C. Accuracy of predicting differentially expressed genes (DEGs) in atopic asthma vs that of the same number of randomly selected genes using the four methods. The list of DEGs was obtained from Forno et al.^20^.

We also examined the prediction accuracy of Methyl-TWAS for eight immune gene pathways available in IMMPORT (**Figure 4, panel B**). Gene expression was predicted most accurately in the chemokine pathway, which regulates airway inflammation in asthma^23^. The second most accurately predicted pathway, antigen presentation, is relevant to dendritic cells^24^ influenced by innate immunity^25^. The third pathway, Natural Killer Cells, modulates immune responses in the airways of subjects with asthma^26^.

The immunity-related genes whose expression was best predicted were those in the human leukocyte antigen (*HLA*) family, such as *HLA-DQB1* (r=0.88), *HLA-DQA1* (r=0.77), and *HLA-DRB5* (r=0.66), known to be associated with asthma and total IgE (**Table 2**). Other genes with well-predicted expression included the *CXCR6/CXCL16* signaling axis (r=0.69), which controls the localization of Resident memory T (T_RM)_ cells to different compartments of the lung and maintains airway T_RM_ cells^27^, and *CD 247* (r=0.66) which is part of the T-cell antigen receptor (TCR) complex and has been associated with moderate to severe asthma ^28^.

**Table 2.** Prediction accuracies (CV-R) of immune genes using four different methods and the pathways the genes belong to.

| Gene ID | Methyl-TWAS | PrediXcan | Hong & Rhee | DNA combi | Immune gene set |
| --- | --- | --- | --- | --- | --- |
| *HLA-DQB1* | 0.883 | 0.83 | 0.879 | 0.895 | Antigen Processing and Presentation |
| *HLA-DQA1* | 0.769 | 0.531 | 0.742 | 0.745 | Antigen Processing and Presentation |
| *CXCR6* | 0.688 | -0.137 | 0.666 | 0.667 | Antimicrobials, Chemokine Receptors, Cytokine Receptors |
| *ZAP70* | 0.675 | 0.034 | 0.655 | 0.641 | Natural Killer Cell, TCR Signaling Pathway |
| *CCL5* | 0.664 | 0.052 | 0.618 | 0.62 | Antimicrobials, Chemokines, Cytokines |
| *HLA-DRB5* | 0.663 | 0.805 | 0.682 | 0.875 | Antigen Processing and Presentation |
| *CD247* | 0.663 | -0.194 | 0.664 | 0.662 | Natural Killer Cell, TCR Signaling Pathway |
| *CD244* | 0.662 | -0.165 | 0.615 | 0.626 | Natural Killer Cell |
| *POMC* | 0.658 | 0.498 | 0.659 | 0.658 | Cytokines |
| *CD48* | 0.651 | -0.167 | 0.644 | 0.647 | Natural Killer Cell |
| *IL1RL1* | 0.651 | -0.094 | 0.561 | 0.536 | Cytokine Receptors, Interleukins Receptors |
| *FCGR3A* | 0.65 | -0.169 | 0.591 | 0.606 | Natural Killer Cell |
| *CD3G* | 0.648 | -0.197 | 0.621 | 0.618 | TCR Signaling Pathway |
| *KLRK1* | 0.646 | 0.123 | 0.496 | 0.468 | Antimicrobials, Natural Killer Cell |
| *CD3D* | 0.639 | -0.172 | 0.626 | 0.621 | TCR Signaling Pathway |
| *SLC40A1* | 0.638 | -0.117 | 0.568 | 0.512 | Antimicrobials |
| *IL12RB1* | 0.635 | -0.014 | 0.624 | 0.627 | Cytokine Receptors, Interleukins Receptors |
| *PRF1* | 0.633 | -0.198 | 0.632 | 0.64 | Natural Killer Cell |
| *PTPRC* | 0.632 | -0.129 | 0.557 | 0.573 | TCR Signaling Pathway |
| *LCK* | 0.629 | -0.161 | 0.604 | 0.59 | Natural Killer Cell, TCR Signaling Pathway |
| *CSF1R* | 0.627 | -0.171 | 0.589 | 0.588 | Cytokine Receptors |
| *ITK* | 0.626 | -0.094 | 0.606 | 0.611 | TCR Signaling Pathway |
| *IL10RA* | 0.624 | -0.009 | 0.616 | 0.621 | Cytokine Receptors, Interleukins Receptors |
| *BPIFA1* | 0.624 | 0.719 | 0.643 | 0.762 | Antimicrobials |
| *SCGB3A1* | 0.623 | -0.187 | 0.528 | 0.473 | Cytokines |
| *CD3E* | 0.623 | -0.192 | 0.626 | 0.617 | TCR Signaling Pathway |
| *CCL26* | 0.621 | -0.128 | 0.567 | 0.496 | Antimicrobials, Chemokines, Cytokines |
| *PIK3R5* | 0.617 | -0.168 | 0.57 | 0.561 | BCR Signaling Pathway, Natural Killer Cell, TCR Signaling Pathway |
| *CCR5* | 0.617 | 0.033 | 0.619 | 0.62 | Antimicrobials, Chemokine Receptors,<br>Cytokine Receptors |
| *CD8A* | 0.617 | 0.03 | 0.606 | 0.582 | Antigen Processing and Presentation,<br>Antimicrobials, TCR Signaling Pathway |

Genes were sorted by Pearson's correlation of Methyl-TWAS

Since EVA-PR includes subjects with and without asthma, we compared the four methods for imputing expression of DEGs in atopic asthma (previously identified in a TWAS conducted in the same cohort^20^). Of 6,058 DEGs, 4,796 were valid genes (r>0.1 for ≥1 of the four methods). Methyl-TWAS -but not PrediXcan-predicted expression of these DEGs significantly better than that of the same number of randomly selected genes (P < 2.2×10^-16^ and 1, respectively) (Figure 4C). Methyl-TWAS also outperformed the other methods in predicting expression of DEGs, suggesting that the combined effects of *cis-* and *trans-*CpGs on expression of DEGs in atopic asthma is much larger than that of SNPs.

### Methyl-TWAS outperforms PrediXcan for *in silico* TWAS of atopic asthma in nasal epithelium

We next conducted *in silico* TWAS of atopic asthma in nasal epithelium using imputed gene expression, performed through fivefold nested cross-validation using EVA-PR data. Methyl-TWAS identified 3,681 (85.2%) of the 4,316 DEGs identified in the previous TWAS using measured expression (FDR-P <0.01)^20^, while PrediXcan could not identify any gene. **Table 3** shows the top 30 TWAS genes (by P value) using observed gene expression. The average CV-r for these 30 genes using Methyl-TWAS was 0.618 while that using PrediXcan was -0.004. These genes include *CST1*^20^ and *ALOX15*^12^. Multiple *ALOX15* SNPs are associated with airway diseases, and DNA methylation and histone modifications in *ALOX15* are associated with airway inflammation^29-32^.

**Table 3.** Comparison of TWAS result using observed gene expression from EVA-PR data, predicted gene expression using Methyl-TWAS, and predicted gene expression using PrediXcan. We used 5-fold nested cross-validation (CV) to measure prediction accuracy (Nested CV-r). The P-values are from TWAS results, empirical Bayes moderated t-statistics using package limma^53^.

|  | Observed<br>gene<br>expression | Predicted gene expression<br>by Methyl-TWAS |  | Predicted gene<br>expression by PrediXcan |  |
| --- | --- | --- | --- | --- | --- |
| Gene ID | P-value | P-value | Nested CV-r | P-value | Nested CV-r |
| *CST1* | 1.70E-32 | 2.50E-17 | 0.69 | 0.32 | -0.11 |
| *TPSAB1* | 3.40E-20 | 7.60E-21 | 0.66 | 0.19 | -0.04 |
| *C3orf70* | 3.70E-20 | 4.10E-21 | 0.72 | 0.81 | 0.15 |
| *CST2* | 1.40E-19 | 9.00E-18 | 0.66 | 0.27 | -0.11 |
| *NTRK2* | 2.90E-19 | 2.20E-18 | 0.71 | 0.94 | -0.096 |
| *CPA3* | 3.40E-19 | 4.70E-24 | 0.69 | 0.35 | -0.04 |
| *ALOX15* | 2.50E-18 | 3.80E-22 | 0.66 | 0.77 | -0.015 |
| *CCL26* | 6.80E-18 | 1.70E-16 | 0.58 | 0.62 | -0.042 |
| *CMYA5* | 1.10E-17 | 1.60E-23 | 0.61 | 0.8 | -0.089 |
| *CISH* | 1.80E-17 | 3.80E-16 | 0.6 | 0.72 | -0.065 |
| *CDH26* | 3.50E-17 | 3.30E-23 | 0.66 | 0.094 | -0.12 |
| *HS3ST4* | 4.10E-17 | 1.90E-16 | 0.59 | 0.6 | -0.032 |
| *TPSB2* | 2.10E-16 | 6.80E-19 | 0.62 | 0.58 | -0.02 |
| *CEP72* | 2.90E-16 | 8.20E-22 | 0.68 | 0.15 | 0.015 |
| *PCSK6* | 3.10E-16 | 4.80E-26 | 0.68 | 0.93 | 0.019 |
| *ADTRP* | 3.40E-16 | 1.20E-19 | 0.65 | 0.52 | -0.066 |
| *GCNT4* | 4.90E-16 | 1.70E-24 | 0.63 | 0.38 | -0.022 |
| *LINC01336* | 2.40E-15 | 2.60E-23 | 0.53 | 0.59 | -0.062 |
| *ANO1* | 2.60E-15 | 1.20E-17 | 0.61 | 0.96 | 0.13 |
| *GSN* | 2.80E-15 | 3.90E-16 | 0.57 | 0.96 | -0.09 |
| *ITLN1* | 2.90E-15 | 6.10E-27 | 0.67 | 0.95 | 0.3 |
| *SLC24A3* | 4.80E-15 | 4.10E-18 | 0.51 | 0.58 | 0.11 |
| *ABO* | 1.20E-14 | 9.40E-17 | 0.57 | 0.24 | 0.37 |
| *WBSCR17* | 1.30E-14 | 2.20E-14 | 0.59 | 0.34 | -0.014 |
| *FETUB* | 1.90E-14 | 2.30E-22 | 0.65 | 0.5 | -0.065 |
| *SLC7A1* | 2.70E-14 | 9.20E-14 | 0.59 | 0.83 | -0.095 |
| *SLC45A4* | 4.90E-14 | 6.20E-21 | 0.54 | 0.81 | -0.007 |
| *DHX35* | 1.00E-13 | 1.80E-13 | 0.53 | 0.73 | -0.011 |
| *RUSC1* | 1.10E-13 | 2.00E-24 | 0.54 | 0.21 | -0.011 |
| *CBX3* | 1.30E-13 | 7.70E-18 | 0.54 | 0.67 | 0.0048 |

*DEG analysis was adjusted for age, sex, plate, and 5 genotypic principal components. Gene was sorted by p-values from observed expression.*

### *In silico* TWAS of atopic asthma in nasal epithelial samples from an external cohort

After building the Methyl-TWAS model trained using EVA-PR data (n=455 subjects; 219 with asthma and 236 controls), we tested this model using public data from an independent cohort (GSE65205, n=69 children, 36 with atopic asthma and 33 non-atopic controls^33^). For prediction accuracy of gene expression, the average r was 0.229 for 10,477 genes (Figure S1), showing a similar range of prediction accuracy in EVA-PR (r=0.256 on average for those genes). We then conducted a TWAS of atopic asthma using imputed gene expression (see Manhattan plot in **Figure 5, Panel A**). Using observed gene expression, we identified 42 DEGs among valid genes (FDR-P <0.05). Using imputed expression, Methyl-TWAS identified 38 (90%) of these 42 DEGs (FDR-P <0.05), all in the same direction of association (**Table 4**). For those 38 DEGs, the average test-r between observed and imputed gene expressions was 0.614.

**Table 4.** Differentially expressed genes (DEGs) in atopic asthma using predicted gene expression trained on EVA-PR data and tested on GSE65205 vs. observed gene expression (GSE65205). Only replicated DEGs were presented. r is the Pearson’s correlation between predicted gene expression and observed gene expression using the test data set.

| Gene ID | Observed expression | | Imputed expression | | $r$ |
| --- | --- | --- | --- | --- | --- |
|  | P-value | FDR | P-value | FDR |  |
| *CCK* | 7.7E-09 | 0.00015 | 3.5E-08 | 0.0000034 | 0.68 |
| *CST1* | 1.5E-08 | 0.00015 | 2.4E-08 | 0.0000027 | 0.89 |
| *CTSC* | 5.3E-08 | 0.00036 | 0.00021 | 0.0013 | 0.58 |
| *CPA3* | 0.00000014 | 0.00059 | 5.6E-08 | 0.0000044 | 0.75 |
| *ANO1* | 0.0000013 | 0.0038 | 0.000068 | 0.00059 | 0.46 |
| *ALOX15* | 0.0000015 | 0.0038 | 0.00000014 | 0.0000084 | 0.82 |
| *NTRK2* | 0.0000017 | 0.0038 | 0.0000055 | 0.0001 | 0.81 |
| *GSN* | 0.0000047 | 0.0088 | 3.8E-09 | 0.00000088 | 0.81 |
| *TPSAB1* | 0.0000054 | 0.0092 | 0.00028 | 0.0017 | 0.57 |
| *ASAP3* | 0.0000078 | 0.012 | 0.00089 | 0.0039 | 0.49 |
| *CXCR6* | 0.0000085 | 0.012 | 0.00024 | 0.0015 | 0.77 |
| *ITLN1* | 0.000016 | 0.018 | 0.00000059 | 0.000023 | 0.62 |
| *CD55* | 0.000017 | 0.018 | 0.0014 | 0.0057 | 0.5 |
| *VWF* | 0.000019 | 0.019 | 3.5E-08 | 0.0000034 | 0.65 |
| *MAP3K4* | 0.000024 | 0.022 | 0.021 | 0.05 | 0.43 |
| *CDH26* | 0.000032 | 0.028 | 0.00000018 | 0.00001 | 0.78 |
| *GJB4* | 0.000035 | 0.028 | 0.00011 | 0.00083 | 0.53 |
| *SLC16A9* | 0.000038 | 0.03 | 0.00036 | 0.002 | 0.69 |
| *HOMER2* | 0.000043 | 0.032 | 0.0017 | 0.0066 | 0.54 |
| *ST6GAL1* | 0.000046 | 0.032 | 0.000063 | 0.00056 | 0.56 |
| *HLA-DMB* | 0.000049 | 0.034 | 0.0000098 | 0.00015 | 0.84 |
| *KLRB1* | 0.000055 | 0.035 | 0.000012 | 0.00018 | 0.75 |
| *EPHA4* | 0.000062 | 0.035 | 0.00023 | 0.0014 | 0.62 |
| *CCL5* | 0.000067 | 0.037 | 0.000011 | 0.00016 | 0.81 |
| *CASP4* | 0.00007 | 0.038 | 0.000016 | 0.00022 | 0.49 |
| *NDC80* | 0.000072 | 0.038 | 0.013 | 0.034 | 0.32 |
| *MARCH3* | 0.000073 | 0.038 | 0.000017 | 0.00022 | 0.53 |
| *SERPINB2* | 0.00008 | 0.039 | 0.0000032 | 0.00007 | 0.56 |
| *ELOVL5* | 0.000084 | 0.04 | 0.000053 | 0.0005 | 0.61 |
| *BDKRB1* | 0.00009 | 0.04 | 0.000027 | 0.00031 | 0.14 |
| *CD6* | 0.0001 | 0.044 | 0.0000032 | 0.00007 | 0.83 |
| *HLA-DOA* | 0.00011 | 0.045 | 0.000023 | 0.00027 | 0.84 |
| *HNRNPA1* | 0.00012 | 0.047 | 0.0078 | 0.022 | 0.17 |
| *SLC2A1* | 0.00012 | 0.047 | 0.00038 | 0.0021 | 0.33 |
| *KAZALD1* | 0.00012 | 0.047 | 0.000031 | 0.00034 | 0.36 |
| *CD3G* | 0.00013 | 0.049 | 0.000062 | 0.00055 | 0.76 |
| *CD74* | 0.00013 | 0.049 | 0.00000085 | 0.000029 | 0.78 |
| *CAPN14* | 0.00014 | 0.05 | 0.0000026 | 0.00006 | 0.68 |

*DEG analysis was adjusted for age, sex, and atopic asthma status. Gene was sorted by p-values from observed expression.*

**Figure 5.**
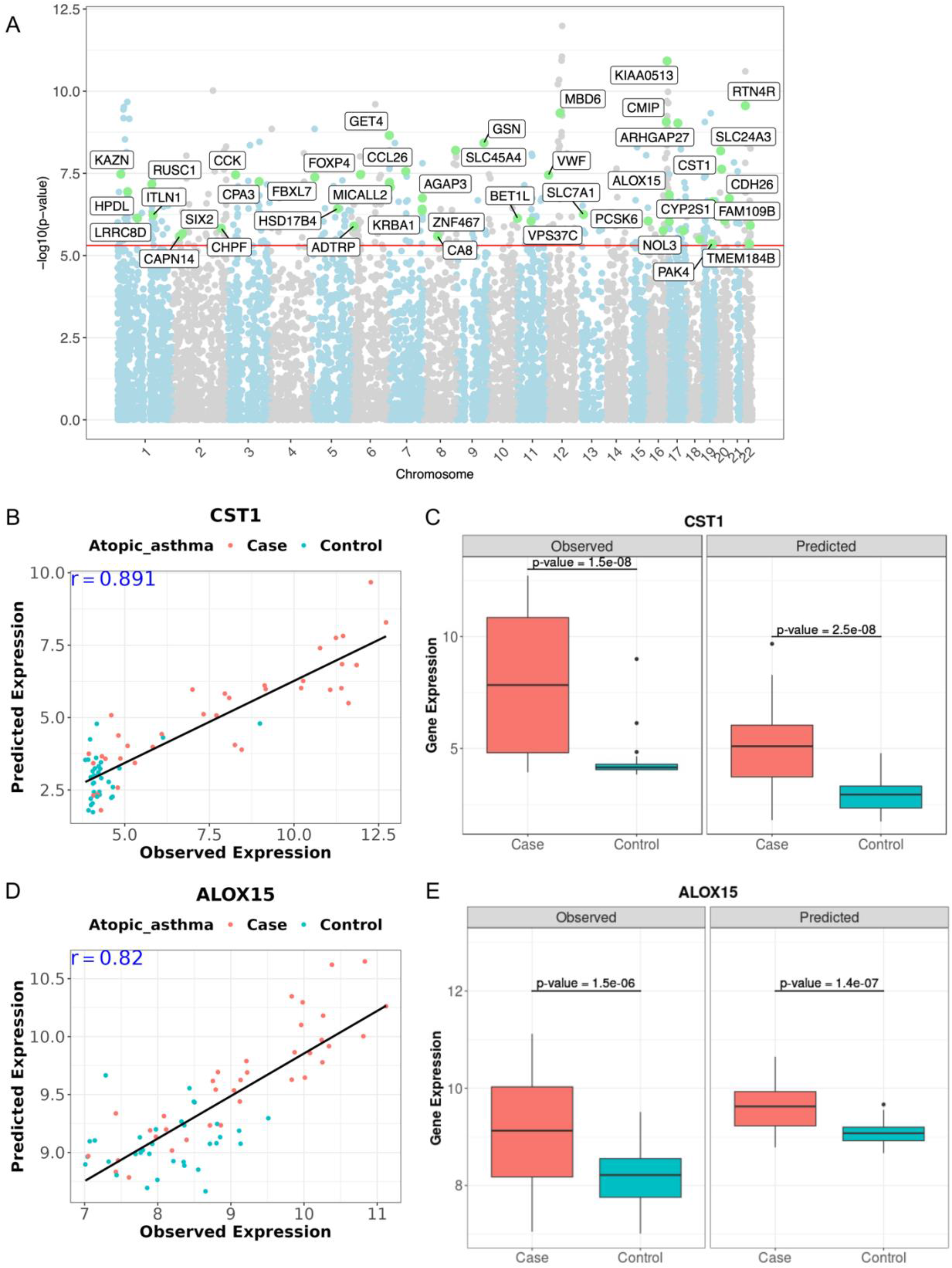
Methyl-TWAS performance of *in silico* TWAS of an external cohort, GSE65205. The models were trained using EVA-PR (N=455); and tested on GSE65205 data (N = 69). A. Manhattan plot of TWAS results using expression levels imputed using Methyl-TWAS. The red line represents the significance threshold after Bonferroni correction (P <0.05). Green dots indicate whether the genes are among the top 200 differentially expressed genes for atopic asthma in a TWAS conducted in EVA-PR^20^. b. Scatter plot of *CST1* expression: observed gene expression of GSE65205 vs. gene expression predicted by Methyl-TWAS. Red dots indicate atopic asthma subjects, and blue dots indicate non-atopic controls. r is Pearson’s correlation between the predicted and observed gene expression levels in the test data set. c. Box plot of *CST1* expression in participants with atopic asthma vs. that in non-atopic controls using observed microarray gene expression (left) and predicted RNA-seq gene expression (right). The P-values are from TWAS results, empirical Bayes moderated t-statistics using package limma^53^. D. Scatter plot of *ALOX15* expression: observed gene expression vs. gene expression predicted by Methyl-TWAS. E. box plot of *ALOX15* expression in atopic asthma vs. that in non-atopic controls using observed (left) and predicted (right) gene expression levels in GSE65205.

The most significant DEG in the *silico* TWAS of atopic asthma that was previously found using observed expression was *CCK* (cholecystokinin), which has been linked to airflow limitation in obesity-related asthma^34^. Other genes included *CTSC* (which influences cell-mediated immune responses and immune cell trafficking in the airways)^35^ and other genes previously discussed. Examples of genes are shown in **Figure 5, panels B-E**. *CST1* is highly predicted by Methyl-TWAS (r=0.89), implying that this gene’s expression is mostly driven by DNA methylation (**Figure 5, panel B**). Using imputed gene expression, Methyl-TWAS accurately identified *CST1* as being upregulated in atopic asthma (**Figure 5, panel C**). A similar pattern was also observed for the imputed expression of *ALOX15* (**Figure 5, panels D-E**).

### Application of Methyl-TWAS to data from white blood cells (WBCs) in EVA-PR

We used Methyl-TWAS to impute gene expression in WBCs from 464 EVA-PR participants with DNA methylation and RNA-seq data. As was the case for nasal epithelium, Methyl-TWAS outperformed PrediXcan (P <2.2×10^-16^; average CV-r were 0.215 and 0.048 for Methyl-TWAS and PrediXcan, respectively, for 8,184 valid genes (r>0.1 for at least one of the four methods) (**Figure 6, panel A**). Using Methyl-TWAS, the prediction accuracy of expression for valid genes in WBCs is similar to that in nasal epithelium (average CV-R was 0.215 in WBCs and 0.246 in nasal tissue for 8,184 valid genes) (**Figure 6, panel A**).

**Figure 6.**
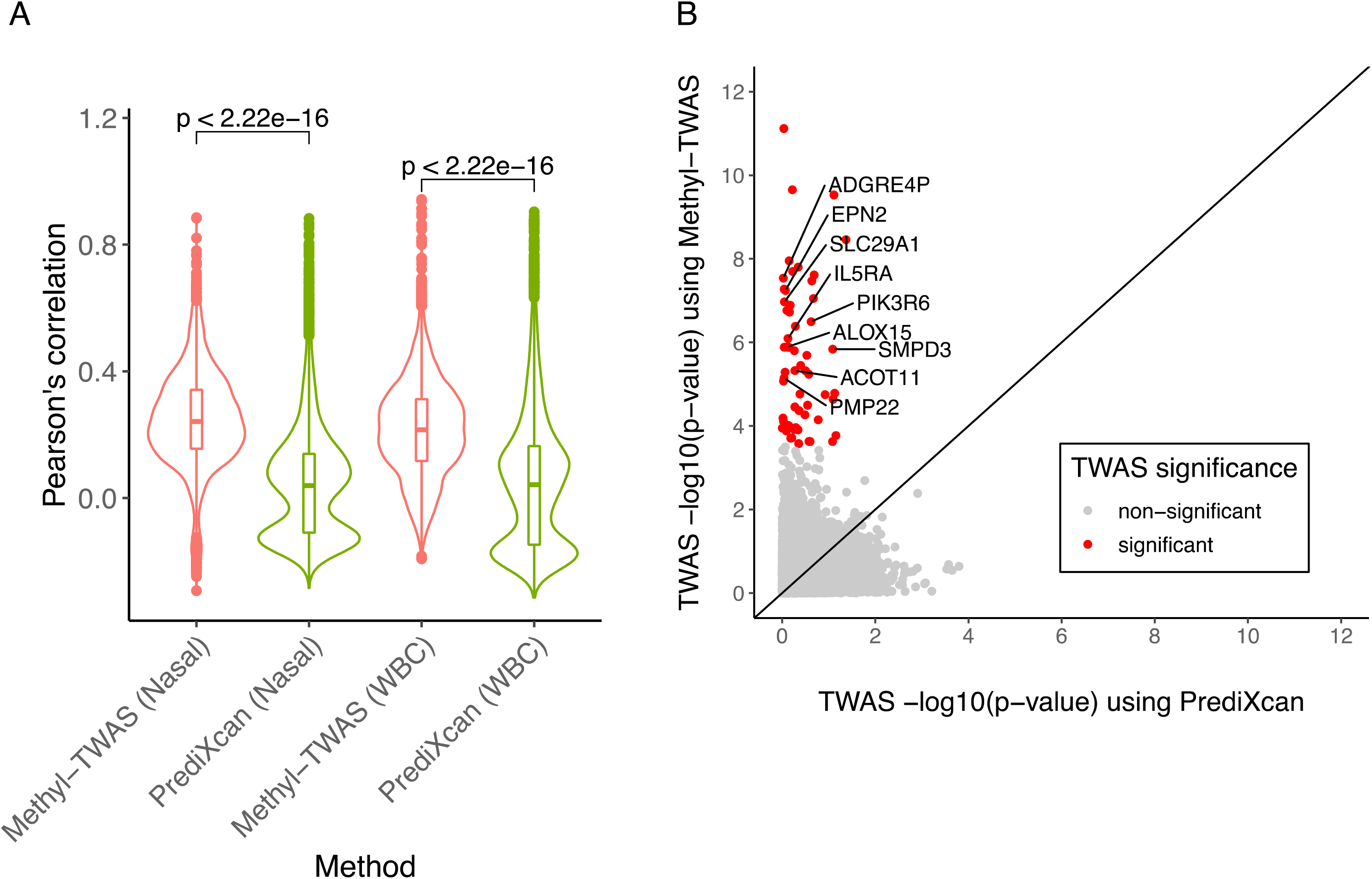
Methyl-TWAS performance in white blood cells from EVA-PR subjects (n=464). A. Comparison of prediction accuracy (cross-validated Pearson’s correlation) between Methyl-TWAS and PrediXcan in nasal epithelium and white blood cells for 8,184 overlapping valid genes between the two types of tissues (CV-r>0.1). A two-sided t-test was used to test the significance of differences. B. Comparison of p-values of TWAS results between Methyl-TWAS and PrediXcan in white blood cells. The red color indicates significant differentially expressed genes (DEGs) for atopic asthma detected by the TWAS analysis (FDR < 0.05). The labeled genes were identified from previous TWAS using observed gene expression data ^36^.

We also conducted *in silico* TWAS of atopic asthma in WBCs. Among 9,139 valid genes, Methyl-TWAS identified 60 DEGs in atopic asthma (FDR-P <0.05), while PrediXcan could not identify any gene (**Figure 6, panel B**). DEGs identified using Methyl-TWAS included nine DEGs reported in the original TWAS using observed expression^36^. Of these nine overlapping DEGs, *SLC29A1* plays a key role in adenosine-regulated suppression of IgE-dependent mast cells^37^, *IL5RA* and *ALOX15* were associated with eosinophil count in a transcriptomic analysis of blood samples from subjects with asthma^38^, and *PIK3R6* and *SMPD3* expression in whole blood was associated with total IgE in subjects with asthma^39^. Other DEGs detected by Methyl-TWAS have been previously implicated in asthma or related traits^39, 40, 41, 42, 43^.

## DISCUSSION

We propose Methyl-TWAS as a novel approach to conduct *in silico* TWAS using long-range CpGs instead of genotype data to improve the accuracy of gene expression imputation and therefore *in silico* TWAS. Methyl-TWAS can estimate epigenetically regulated/associated expression (eGReX), which includes parts of genetically (GReX) and environmentally regulated expression, trait-altered expression, and tissue-specific expression, since DNA methylation is affected by all these ^44-46^ (**Figure 1**). Note that traditional TWAS methods such as PrediXcan can only estimate GReX.

In the first step of predicting gene expression, using nasal epithelium from youth with asthma and control subjects, Methyl-TWAS outperformed three methods, especially PrediXcan. The predictive power of *cis-*methylation (Hong and Rhee’s approach) on gene expression was much higher than that of *cis-*SNPs (PrediXcan). This shows that the proportion of nasal epithelial gene expression explained by epigenetics is much greater than that explained by genetics for most genes. Further, the prediction accuracy of long-range DNA methylation (Methyl-TWAS) was higher than that of *cis*-range DNA methylation (Hong and Rhee’s approach), and this was not improved by adding SNPs to DNA methylation data (DNA combi) for most genes. This implies that for SNPs and CpGs associated with gene expression, DNA methylation mediates a substantial proportion of the effect of SNPs on nasal epithelial gene expression (∼89% in our previous study of asthma-susceptibility SNPs)^47^. Further, Methyl-TWAS predicts expression of a substantial proportion of DEGs of atopic asthma and genes in immune-related pathways while PrediXcan does not.

When we performed *in silico* TWAS, Methyl-TWAS markedly outperformed PrediXcan in identifying DEGs in atopic asthma. Moreover, Methyl-TWAS outperformed PrediXcan in an *in silico* TWAS of atopic asthma in a tissue other than nasal epithelium (WBCs), with DEGs implicated in asthma and total IgE. The summary of a comparison of the four methods is shown in **Table S3**.

Methyl-TWAS can be used if both DNA methylation and gene expression data are available for any tissue. Indeed, we found good prediction accuracy for gene expression and *in silico* TWAS in both nasal epithelium and WBCs (Figure 6A, 6B). For studies of nasal epithelium, we already trained the model, and thus users need only methylation data and a phenotype to impute gene expression and conduct *in silico* TWAS result using Methyl-TWAS. Indeed, we found similar prediction accuracy for gene expression in nasal epithelium from participants in GSE65205, which, like EVA-PR, included individuals with and without asthma (**Figures 5)**.

While R^2^ is often used to assess the prediction accuracy of gene expression for *in silico* TWAS, this measure can inflate such accuracy, as 45.43% of imputed expression levels using PrediXcan were negatively correlated with observed gene expression in the current analysis. Thus, using Pearson’s correlation or other measure that shows the direction of a correlation is preferable to using R^2^ for *in silico* TWAS.

We recognize several study limitations. First, we could not conduct a simulation study using both genotypes and CpGs because their correlation is largely unknown and because of the challenges posed by trying to generate an ultra-high dimensional covariance matrix for the simulation study (tens of millions of SNPs times 450K DNA methylation CpGs/2). Second, there is no dataset with genome-wide genotypes, CpGs, and gene expression in nasal epithelium within the same cohort other than EVA-PR, limiting comparisons between PrediXcan and Methyl-TWAS in a test data set. Third, we do not know the directionality of associations when using CpGs to impute gene expression in Methyl-TWAS: in some cases, DNA methylation affects gene expression and in others, gene expression affects DNA methylation. However, identifying the causality of DNA methylation on gene expression is beyond the scope of the current analysis.

By incorporating long-range DNA methylation, Methyl-TWAS provides significantly improved prediction accuracy of gene expression compared to traditional genotype-based methods, which in turn facilitates more accurate identification of DEGs when conducting *in silico* TWAS. Through a user-friendly R package, Methyl-TWAS allows researchers to conduct accurate *in silico* TWAS using a growing body of publicly available DNA methylation data to study various diseases, including asthma and other immune-related diseases.

## Acknowledgements

This work was supported by grants HL079966, HL117191, MD011764, and HL119952 (to J.C.C.), U54 MD007587 (to E.A-P. and G.C.), and K01 HL153792 (to S.K.) from the U.S. National Institutes of Health (NIH). The contribution of E.F. was supported by NIH grant Hl149693, and that of W.C. was supported by NIH grant HL150431 and NSF grant 2225775. Data analysis was supported in part by the HTC cluster of the University of Pittsburgh Center for Research Computing, which is supported by NIH award number S10OD028483.

## Author contributions

S.K. designed and conducted experiments and wrote the paper; Y.Q. conducted the experiments; H.P. E.F. W.C. and J.C. provided helpful comments; M.Y. and Z.X. pre-processed other validation datasets; J.C. and W.C. edited the manuscript; J.C. collected and provided the EVA-PR dataset.

## Data and code availability

Genome-wide DNA methylation data and gene expression array data from the replication cohort were deposited in the Gene Expression Omnibus (GEO accession No: GSE65205). The estimated coefficients for each CpG for each gene using EVA-PR data trained by Methyl-TWAS are available in GitHub: https://github.com/YidiQin/MethylTWAS/tree/9c990c32161a6688c7bd33825b6e3554490c7afd/data/test.10M.EVA_PR.cvfit.Rdata

We have no consent for public sharing of individual-level omics data from participants in EVA-PR, who are members of a vulnerable population. Qualified investigator should contact the corresponding author regarding requirements and processes for data sharing.

R-package Methyl-TWAS is downloadable from https://github.com/YidiQin/MethylTWAS.

