## Supplementary material for "Methyl-TWAS: A powerful method for *in silico* transcriptome-wide association studies (TWAS) using long-range DNA methylation": Table S3

**Table S3. Comparison of four methods for gene expression imputation and/or *in-silico* TWAS**

| | Methyl-TWAS | PrediXcan | Hong and Rhee | DNA combi |
|---|---|---|---|---|
| Data | CpGs $\pm 10Mb$ from promoter region of each gene | SNPs $\pm 1Mb$ from promoter region of each gene | CpGs $\pm 1Mb$ from promoter region of each gene | DNA SNP and CpGs $\pm 10Mb$ from Promoter region of each gene |
| Ability to conduct gene expression imputation | Yes | Yes | Yes | Yes |
| Ability to conduct in silico-TWAS | Yes | Yes | Not applicable | Not applicable |
| Gene expression prediction accuracy (ranking of average accuracy across genes) | 1 | 4 | 3 | 2 |
| Genes whose expression is better predicted | Immunity-related genes/differentially expressed genes (DEGs) for atopic asthma | Genes in metabolic and biosynthetic pathways | Immunity-related genes/DEGs for atopic asthma | Immunity-related genes |
| In Silico-TWAS results | Can identify most DEGs for atopic asthma | Cannot identify any DEG for atopic asthma | Not applicable | Not applicable |
